# Long-range inhibitory neurons mediate cortical neurovascular coupling

**DOI:** 10.1101/2022.10.11.511811

**Authors:** Catherine F. Ruff, Fernanda Juarez Anaya, Samuel J. Dienel, Adiya Rakymzhan, Alain Altamirano-Espinoza, Jay Couey, Alan M. Watson, Kenneth N. Fish, Bryan M. Hooks, Maria E. Rubio, Aihua Su, Sarah E. Ross, Alberto L. Vazquez

## Abstract

Neuronal activity evokes a vascular response that is essential to sustain brain function. We show that neurovascular coupling (NVC) is mediated by long-range projecting GABAergic neurons that express Tacr1. Whisker stimulation elicited Tacr1 neuron activity in the barrel cortex through feed-forward excitatory pathways. Optogenetic activation of Tacr1 neurons elicited vasodilation, whereas inhibition significantly reduced whisker-evoked hemodynamic responses. Moreover, vasodilation was preceded by capillary pericyte activity, demonstrating a mechanism for NVC.

## Main Text (Introduction, Results, Discussion)

Neurovascular coupling (NVC) is the mechanism that adjusts cerebral blood flow (CBF) to match local neural activity^1^. This coupling mechanism is the basis of functional human imaging studies of brain function (i.e., BOLD fMRI). Furthermore, dysregulation of NVC is thought to contribute to neurodegenerative diseases such as Alzheimer’s Disease^2-5^. However, the neuronal basis for NVC remains unclear.

Cortical GABAergic neurons that express neuronal nitric oxide synthase (nNOS or Nos1) have been postulated to play a role in CBF because they release nitric oxide (NO), a gaseous signaling molecule that induces rapid vasodilation^6-8^. Recent single-cell sequencing studies have suggested that Nos1 is expressed in a well-defined transcriptomic subset of GABAergic neurons^9, 10^. Through multiplex fluorescent in situ hybridization (FISH) we identified and characterized this population in mouse cortex as a somatostatin-expressing subpopulation that shows an almost complete overlap of three markers: *Nos1, Tacr1*, and *Chodl* (Fig. 1a-c and Extended Data Fig. 1a,b,d,e), consistent with sequencing studies. In human cortex, we likewise observed that a small subpopulation of *SST-*positive neurons co-expressed *NOS1, TACR1* and *CHODL*, and that these neurons were sparse in superficial cortical layers and more concentrated in layer 6 and white matter (Fig. 1d,e and Extended Data Fig. 1c,f-h). These findings are consistent with previous descriptions of Type 1 nNOS neurons in guinea pig, rat and monkey, and suggest conservation of this cell type across species^11^.

**Figure 1.**
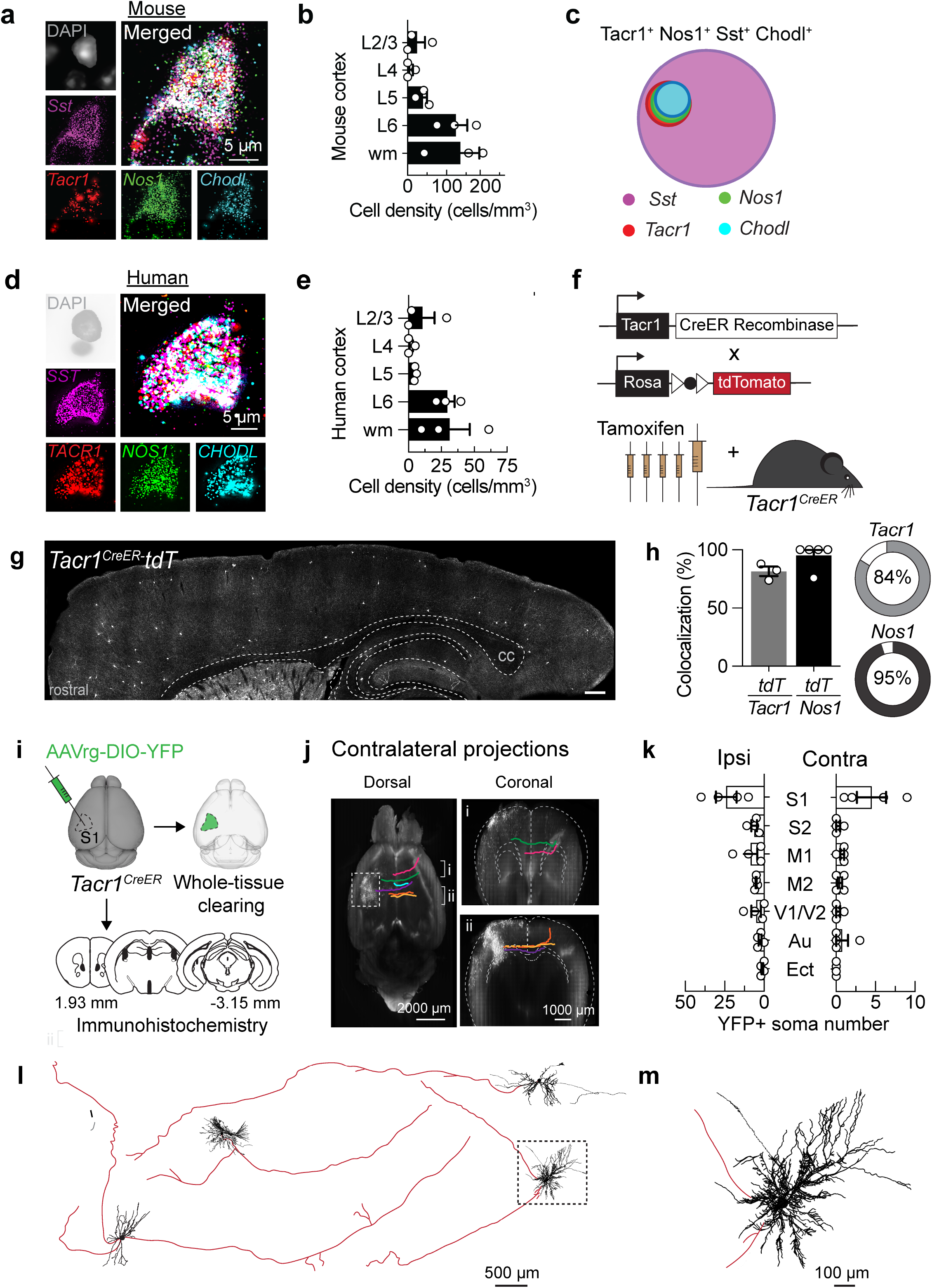
Tacr1 marks a conserved population of long-range GABAergic neurons. Representative multiplex fluorescent in situ hybridization (FISH) images demonstrating colocalization of *Sst/SST* (purple), *Tacr1/TACR1* (red), *Nos1/NOS1* (green) and *Chodl/CHODL* (blue) in cortical neurons in C57BL/6 mouse (**a**) and human (**d**). Quantification of neuronal density of quadruple-labeled cells in cortex and white mater in mouse (**b**, n = 3 mice) and human (**e**, n = 3 subjects) White matter, wm. **c**, Venn diagram depicting the intersectionality of these neuron populations in mouse. **f**, Cartoon depicting strategy to target *Tacr1*^*CreER*^ neurons. **g**, Representative sagittal section of *Tacr1*^*CreER*^-mediated tdTomato (tdT) expression in the cortex after tamoxifen administration. Dashed lines are approximate anatomical borders. Corpus callosum, cc. **h**, Quantification of the percent *Tacr1*- or *Nos1*-expressing neurons that expressed tdT (specificity) and the percent of tdT^+^ neurons that co-expressed *Tacr1* or *Nos1* (recombination efficiency). **i**, Schematic depicting experimental design. AAV-retrograde(rg) was injected into mouse somatosensory cortex. Brains were processed for whole-tissue clearing or immunohistochemistry. **j**, Representative contralateral projections of *Tacr1* neuron traced in cleared brains (n = 2 *Tacr1*^*CreER*^ mice). **k**, Quantification of Tacr1 neuron cell bodies in various brain regions following the injection AAVrg-DIO-YFP in *Tacr1*^*CreER*^ mice (n = 4). **l**, Representative neuronal reconstructions of Td-tomato labeled *Tacr1* neurons from cleared brains showing long-range axons (red) and dendrites (black). **m**, Inset from l to illustrate Tacr1 dendrites. Data are mean ± s.e.m and dots represent data points from individual mice or subjects.

Cortical GABAergic neurons are generally considered to be interneurons, but recent evidence suggests that Type 1 nNOS neurons may extend long-range (>1.5 mm) axonal projections^12, 13^. To visualize these neurons and examine their vasoregulatory role in more detail, we generated a *Tacr1*^*CreER*^ knock-in allele, thereby targeting Type 1 nNOS neurons without disrupting the Nos1 locus (Fig. 1f-g). This allele faithfully recapitulated the endogenous expression of Tacr1 at both the level of protein and mRNA (Fig. 1g-h, Extended Data Fig 2a-e and 3a-g), and successfully targeted the majority of neurons that express Nos1 at high levels (Fig. 1h, Extended Data Fig. 2f and 3e,g)^14^. To visualize the morphology of these neurons, we performed Cre-dependent retrograde viral labeling in somatosensory cortex of *Tacr1*^*CreER*^ mice combined with either immunohistochemistry or whole-tissue clearing and ribbon scanning confocal microscopy (Fig. 1i). Labeled soma were widely distributed across the cortex, consistent with the idea that Tacr1 neurons have long-range projections that extend within and across hemispheres (Fig. 1j,k and Extended Data Fig. 4a-c). Moreover, interhemispheric projections from Tacr1 neurons dominated homotopic S1, paralleling known pyramidal cortico-cortical connections in the contralateral cortex (Fig. 1k). Single-cell tracing revealed that dendrites from Tacr1 neurons have extensive radial branches extending ∼500 μm from the soma, whereas axons from Tacr1 neurons had limited branching, and extended several millimeters (Fig. 1m,l and Extended Data Fig. 4f-g).

To begin investigating the role of Tacr1 neurons in NVC, we examined whether these neurons respond to sensory stimulation using two-photon microscopy in awake mice expressing GCaMP6f (Fig 2a). Whisker stimulation triggered a rapid and robust Ca^2+^ transient in Tacr1 neurons in the contralateral barrel cortex (Fig. 2b-d) that was similar in dynamics to those observed in pyramidal neurons (Fig. 2b, inset). To investigate the synaptic basis of this evoked activity, we performed subcellular ChR2-assisted circuit mapping (sCRACM), expressing ChR2 in either pyramidal neurons or thalamic neurons and recording from tdTomato(*tdT*)-expressing *Tacr1*^*CreER*^ neurons (Fig. 2e). In either case, optical simulation in the presence of TTX and 4-AP evoked post-synaptic currents in Tacr1 neurons, consistent with monosynaptic cortico-cortical and thalamo-cortical excitatory input (Fig. 2f-g). Thus, Tacr1 neurons integrate local activity through feed-forward excitatory pathways from cortex and thalamus.

**Figure 2.**
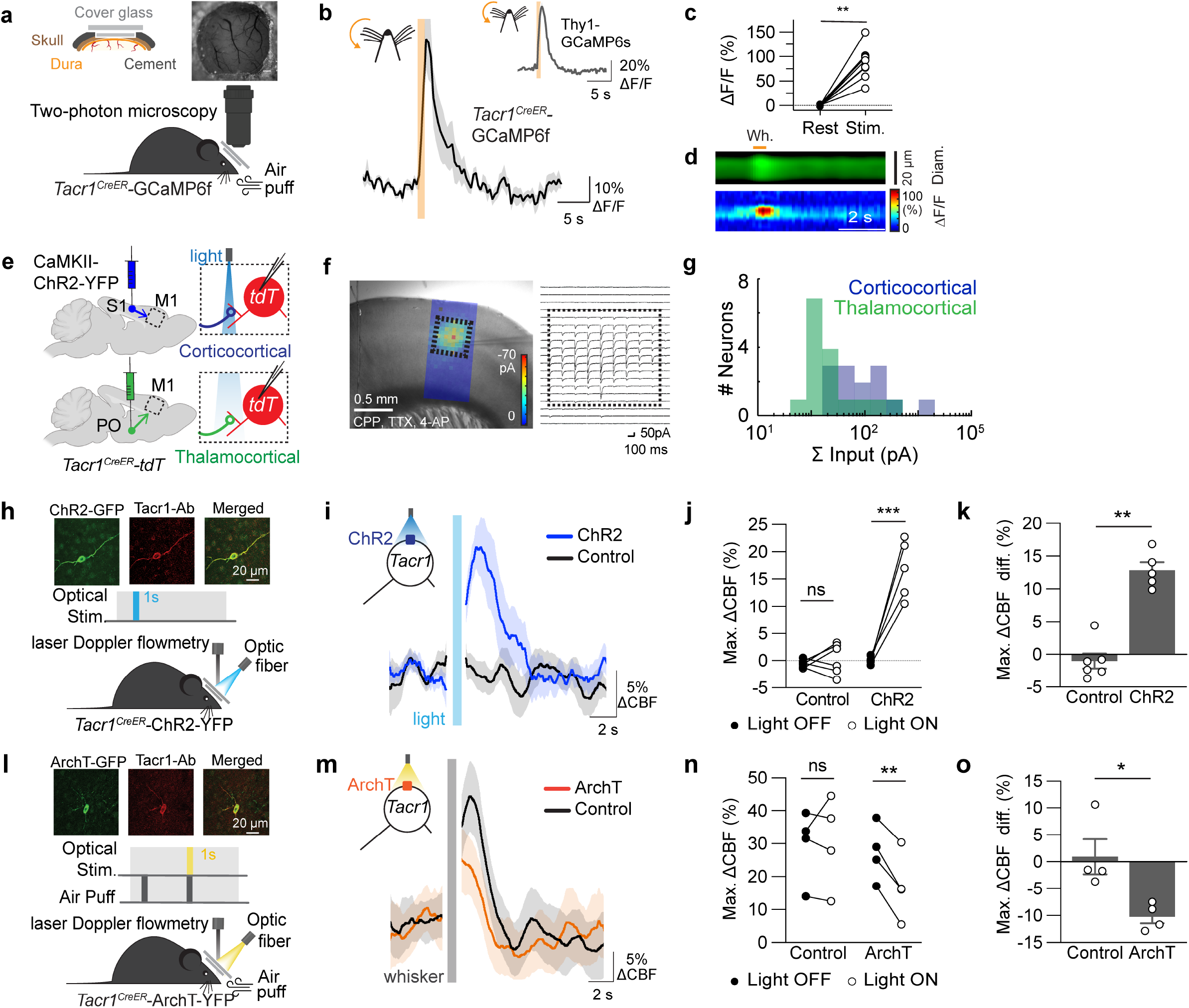
Tacr1 neurons are recruited for sensory-evoked NVC and are required for NVC. **a**, Top: Schematic of chronic cranial window design and representative macroscopic image of a cranial window in mouse. Bottom: Schematic depicting Ca^2+^ imaging by two-photon microscopy in somatosensory (S1) cortex. Sensory stimulation (air puff) to the contralateral whisker pad was performed to evoke a hemodynamic response. **b**, Time course of the percent change in Ca^2+^ signal (ΔF/F) in mice expressing GCaMP6 in *Tacr1* neurons or *Thy1* (pyramidal) neurons (inset). **c**, Maximum percent change in Ca^2+^ signal in mice expressing GCaMP6f in *Tacr1*^*CreER*^ neurons before and after whisker stimulation (\*\**P* = 0.0039, n = 9 neurons in 5 mice). **d**, Representative line profiles showing ΔF/F from a GCaMP6f-expressing *Tacr1* neurons and ΔD/D of a nearby vessel. Orange bar represents whisker stimulation (1 s, 50 Hz, 50 ms). **e**, Schematic of subcellular ChR2-assisted circuit mapping (sCRACM) showing optogenetic stimulation of ChR2-expressing cortical or thalamic excitatory neurons and simultaneous recording from tdT-expressing *Tacr1*^*CreER*^ neurons in M1. **f**, Left: Example sCRACM map superimposed on a brightfield image of a M1 brain slice. Right, example of averaged excitatory postsynaptic currents (EPSC_sCRACM_) recorded from a *Tacr1*^*CreER*^ neuron displayed on the grid corresponding to the light stimulus location. **g**, Histogram of synaptic strenght for cortical (S1, green; n=13) and thalamic (PO, blue; n=16) input to *Tacr1*^*CreER*^ neurons in M1. **h-k**, Acute optogenetic excitation of cortical ChR2-expressing *Tacr1* neurons. **h**, Top: IHC showing colocalization of ChR2 (green) and Tacr1 (red) protein. Middle: Optogenetic stimulation protocol (1 s, 5 Hz, 30 ms pulse width, blue light). Bottom: Experimental setup demonstrating continuous cerebral blood flow measurement by laser Doppler flowmetry (LDF) during optical excitation in awake, head-fixed mice. Time course of the change in CBF (**i**), maximum change in CBF (**j**) and percent change in CBF from light OFF (**k**) in mice expressing ChR2 in *Tacr1* neurons (n=5 mice, 10 trials per mouse) compared to control mice (n = 6, 10 trials per mouse). Activation of cortical *Tacr1* neurons significantly increased CBF relative to light OFF (**j**, ***P = 0.0004) and compared to control mice (**k**, **P = 0.0043). **l-o**, Acute optogenetic silencing of cortical ArchT-expressing *Tacr1* neurons. **l**, Top: IHC showing colocalization of ArchT (green) and Tacr1 (red) protein. Middle: Optogenetic silencing protocol (1 s, 5 Hz, 5 ms pulse width; yellow light). Bottom: Experimental setup demonstrating continuous CBF recording by LDF during optical inhibition in the presence of sensory stimulation (air puff; 1 s, 10 Hz, 50 psi) in awake, head-fixed mice. Time course of the change in CBF (**m**), maximum change in CBF (**n**) and percent change in CBF from to light OFF (**o**) in mice expressing ArchT in *Tacr1* neurons compared to control mice (n=4 mice, 10 trials per mouse). Inhibition of cortical *Tacr1* neurons significantly decreased sensory-evoked CBF relative to light OFF (**o**, \**P* =0.0038) and compared to control mice (**n**, \*\**P* = 0.0143). Data are mean ± s.e.m. Statistical significance was determined by paired, nonparametric two-tailed *t*-test (d), paired, parametric, two-tailed *t*-test (j, n) or unpaired, nonparametric two tailed *t*-test (d, k, o). ns, not significant. Error bars and shaded areas are s.e.m.

To test whether the activation of Tacr1 neurons is sufficient for vasodilation, we optogenetically activated *Tacr1*^*CreER*^ neurons and measured blood flow *in vivo* using laser Doppler flowmetry (Fig 2h and Extended Data Fig. 5a-d). Although Tacr1 neurons are sparse, representing approximately 1% of cortical neurons, optogenetic activation of Tacr1 neurons significantly increased CBF with a magnitude that was comparable to that observed upon whisker stimulation (Fig. 2i-k and Extended Data Fig. 5g). In contrast, blue light stimulation did not elicit a blood flow response in ChR2-negative animals (Extended Data Fig. 5h-n). To address the necessity of Tacr1 neurons for NVC, we tested whether inhibition of *Tacr1*^*CreER*^ neurons in S1 was sufficient to reduce the NVC response to whisker stimulation (Fig 2l). Indeed, the whisker-evoked NVC response was significantly reduced upon light-mediated inhibition of archaerhodopsin (ArchT)-expressing Tacr1 neurons (Fig. 2m-o and Extended Data Fig. 6a-c). In contrast, yellow light stimulation in control mice did not in itself elicit a CBF response, and had no significant effect on the whisker-evoked CBF response (Extended Data Fig. 6d-f). Taken together, these experiments indicate that Tacr1 neurons are necessary and sufficient for NVC.

An unresolved question is whether NVC is initiated by the relaxation of vascular smooth muscle cells (VSMCs) or capillary pericytes^15-18^. To investigate the site of NVC within the neurovascular tree, we monitored mural cell activity and cortical blood vessel diameter using two-photon microscopy through cortical windows in awake, head-fixed mice. (Fig. 3a-c). ChR2 was expressed in *Tacr1*^*CreER*^ neurons, GCaMP6f was expressed in mural cells (VSMCs and pericytes), and the vasculature was visualized with an intravascular dye (Fig. 3a-c). We found that optogenetic activation of cortical Tacr1 neurons resulted in a transient decrease in cytosolic calcium [Ca^2+^]_i_ in both pericytes and VSMCs, consistent with relaxation (Fig 3d,e). This relaxation was accompanied by vasodilation of both capillaries (1^st^-4^th^ order branches) and arterioles (Fig 3d,f). Interestingly, the latency to relaxation was significantly shorter in pericytes compared to VSMCs (Fig 3g). Complementary to these findings, the latency to vessel dilation was significantly shorter in capillaries compared to arterioles (Fig 3h). Similar results—with the decrease in [Ca^2+^]_i_ in pericytes preceding the decrease in [Ca^2+^]_i_ VSMCs, and changes increase in capillaries diameter preceding the increase in arteriolar diameter—were observed when vasodilation was triggered by whisker stimulation (Extended Data Fig. 7 a-h). These findings support the idea that blood flow observed in early capillaries (>4^th^ order) is actively regulated by contractile pericytes, which ensheath initial capillary branch points, and suggest that NVC is initiated at these capillaries rather than arterioles.

**Figure 3.**
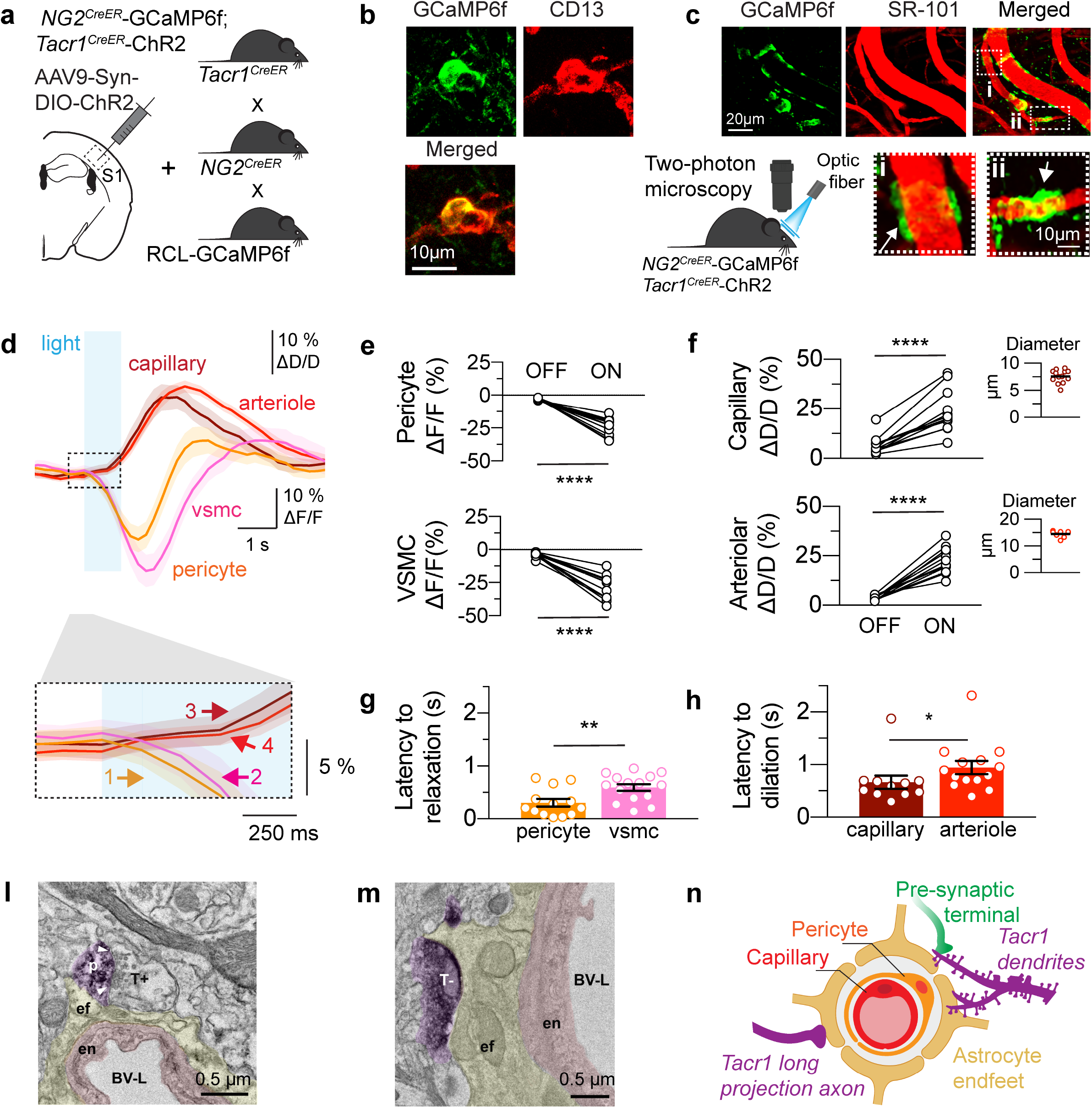
Tacr1-neuron evoked NVC is mediated by contractile capillary pericytes. **a**, Schematic depicting strategy to express ChR2 in *Tacr1*^*CreER*^ neurons and GCaMP6f in *NG2*^*CreER*^ mural cells (Note, with this strategy, GCaMP6f is also expressed in *Tacr1*^*CreER*^ neurons, although it is not relevant in these experiments). **b**, White arrows indicate capillary pericytes that express GCaMP6f (green) and CD13 (pericyte marker, red). **c**, Representative *in vivo* two-photon image (maximum intensity projection) showing GCaMP6f-expressing VSMCs (green) and SR-101 labeled vasculature (red). In merged image, white dashed boxes shown enlarged at bottom. Bottom: VSMC (white arrow) on arteriole (i) and ensheathing pericyte (white arrowhead) on capillary (ii). Schematic of light stimulation and simultaneous recording of Ca^2+^ in VSMCs/pericyte and vessel diameter. **d**-**h**, VSMCs/pericyte and vascular dynamics upon optogenetic stimulation in *Tacr1* neurons (n = 5 mice). **d**, Top: Time course of the change in VSMC/pericyte Ca^2+^ signal and subsequent vessel diameter. Area outlined by black dashed rectangle shown at bottom. Bottom: VSMC/pericyte relaxation (Ca^2+^ decrease) precedes vascular dilation during light stimulation. Numbers indicate response onset to light stimulation; colors correspond to the cell type shown above. **e**, Maximum percent change in pericyte (top) and VSMCs (bottom) ΔF/F, or Ca^2+^ signal (\*\*\*\**P* < 0.0001, 12 pericytes and 13 VSMCs). **f**, Maximum percent change in capillary (top) and arteriolar (bottom) ΔD/D, diameter (\*\*\*\**P* < 0.0001, 12 capillaries and 12 arterioles). Insets show resting vessel diameter. **g-h**, Latency to onset of mural cell relaxation (\*\**P* =0.0093) or vascular dilation (\**P* = 0.0128) upon optogenetic stimulation. **l-m**, Electron microscopy of *Tacr1*^*CreER*^ (purple) processes to the microcirculation. Blood vessel (BV, pink), includes endothelial cell and pericyte. **l**, Representative image of *Tacr1*^*CreER*^ dendritic spine (purple) forming an asymmetrical synapse with a putative excitatory terminal (T+). White arrowheads point out the edges of the postsynaptic density (p). *Tacr1*^*CreER*^ distal dendrite receiving a putative excitatory terminal (T+) also contact perivascular astrocyte endfoot (ef, yellow), forming a suspected neuronal-astrocytic-vascular tripartite functional unit. **m**, *Tacr1*^*CreER*^ putative inhibitory axon terminal (T-) contacting astrocytic endfeet. en: endothelial cell; BV-L: blood vessel-lumen **n**, Schematic of the neurovascular unit including proposed perivascular location of *Tacr1* processes. Data are mean ± s.e.m; statistical significance was determined by paired, two-tailed *t*-test (**e**,**f**) unpaired, two-tailed Mann-Whitney test (**g**,**h**,**j**,**k**).

To visualize the ultrastructure of Tacr1 neurons in the context of the neurovascular unit and the possibility of contacts between Tacr1 processes and vascular elements, we used transmission electron microscopy of immunolabeled *Tacr1*^*CreER*^ neurons that expressed ChR2. We found that both presynaptic and postsynaptic terminals of Tacr1 neurons make contacts with astrocytes and/or astrocytic end-feet that surround blood vessels (Fig. 3l,m and Extended Data Fig. 8a-e). Moreover, we observed juxtaposition of an astrocytic end-foot to the asymmetrical synapse between a putative excitatory presynaptic terminal and Tacr1 postsynaptic terminal (Fig. 3l). These results show that both the presynaptic (axonal) terminals and the postsynaptic (dendritic) processes of Tacr1 neurons are intimately associated with the neurovascular unit, and thus anatomically positioned to mediate NVC (Fig. 3n).

In summary, we found that Tacr1 neurons are a distinct GABAergic neural population co-expressing Sst, Nos1, and Chodl in both mouse and human cortex. Our study revealed that Tacr1 neurons are necessary and sufficient for feed-forward increases in cerebral blood flow in the barrel cortex that are recruited by whisker stimulation. Activation of Tacr1 neurons caused the relaxation of pericytes and VSMCs, as evidenced by a decrease in [Ca^2+^]_i_ in mural cells that was accompanied by the vasodilation of arterioles and capillaries. Moreover, our data sheds light on the controversial question of where vasodilation is initiated by providing evidence that contractile pericytes at the arteriolar-capillary junction are the first to respond *in vivo*, consistent with previous studies^19^. Finally, our anatomical data identified perivascular Tacr1 postsynaptic dendrites and presynaptic axonal terminals that are well within range to signal to the neurovascular unit via vasoactive substance (e.g., NO) or other neurotransmitters (e.g., GABA). The novel finding that Tacr1 neurons form a tripartite synapse is consistent with previous observations indicating that glutamate triggers the release of NO, and is consistent with a direct NO-mediated mechanism for mural cell relaxation and subsequent vasodilation. Moreover, these findings raise the intriguing possibility that NVC might occur within subdomains of the Tacr1 dendritic tree, thereby enabling highly localized control of blood flow. Given the association between impaired NVC and neurodegenerative diseases, it will be important to investigate whether dysfunction in Tacr1 neurons contributes to disease pathogenesis.

## Methods

### Experimental animals

All animals were cared for in compliance with the National Institutes of Health guidelines and approved by the University of Pittsburgh Institutional Animal Care and Use Committee. Previously generated^14^ *Tacr1*^*CreER*^ mice (also called *NK1R*^*CreER*^ mice) were bred in house and maintained on a C57BL/6 background. The following mice strains were used: wild type (C57BL/6J, Charles River, no. 027), *NG2*^*CreER*^ (ref. ^20^, JAX, no. 008538), Ai9 (JAX, no. 007909), Ai32 (JAX, no. 012569), Ai35 (ref. ^21^; JAX, no. 012735), Ai95 (JAX, no. 028865), Thy1-GCaMP6s (JAX, no. 024275). All mice were maintained on a C57BL/6J background and both males and females were used. For mice expressing *CreER*, tamoxifen (Sigma-Aldrich, T5648) was dissolved in corn oil at a concentration of 20 mg ml^−1^ and injected into peritoneal cavities at a dose 75 mg kg^-1^ for 5 consecutive days. Animals recovered for at least one week following the last tamoxifen treatment before cranial surgery or dissections were performed. Sample sizes were determined by a power calculation based on previous pilot data and representative sample sizes from previous literature that had similar experiments.

### Immunohistochemistry

Tamoxifen-treated *Tacr1*^*CreER*^; *Rosa26*^*lsl-tdT*^(Ai9) adult mice were transcardially perfused with 4% paraformaldehyde. Brains were dissected out of the skull and post-fixed overnight. Sagittal or transverse 40 μm sections were cut on a vibratome (Leica 1200) and processed for free-floating immunohistochemistry. Brain sections were blocked with 10% goat or donkey serum, phosphate-buffered saline + triton (0.1% Triton X-100, PBST) and stained overnight at 4 °C with the following primary antibodies at the indicated concentrations: GFP (1:1000; ThermoFisher A-11122; Aves Lab Inc., GFP-1020), Tacr1 (1:10,000; Sigma S8305), nNOS (1:2000, Abcam Ab1376), neuropeptide Y (1:1000; Peninsula Laboratories, LLC; T-4070), parvalbumin (1:2000; SWANT; PVG-213), RFP (1:1000; Rockland; 600-401-379S) and somatostatin (Peninsula Laboratories, LLC; T-4547), followed by the corresponding Alexa Fluor-conjugated secondary antibodies (1:500, ThermoFisher). Sections were mounted in Fluoromount-G (SouthernBiotech) and imaged with confocal microscopy (Nikon A1R) and fluorescent microscopy (Nikon 90i). Anatomical regions were identified using the Paxinos and Allen Institute Mouse Brain Atlases.

### Transmission electron microscopy

Adult *Tacr1*^*CreER*^; *Rosa26*^*lsl-ChR2-eYPF*^ mice were anesthetized and then transcardially perfused with 0.1M phosphate buffer (PB) pH=7.2, followed by 4% PFA and 0.5% glutaraldehyde in 0.1 M phosphate buffer (PB). Brains were post-fixed for 90 minutes in the same fixative at 4ºC. Following fixation, 80 μm coronal sections were cut by vibratome in 0.1M PB and then cryoprotected with an ascendant gradient of glycerol in 0.1M PB. The glycerol gradient involved two 30-minute incubations (10% and 20%), followed by an overnight incubation in 30% glycerol. Sections were frozen on dry ice and thawed in 0.1M PB. For the immunohistochemistry, sections were blocked with 10% normal goat serum for 1 hour at room temperature (RT), incubated overnight at 4ºC with a rabbit anti-GFP primary antibody (1:1000; ThermoFisher A-11122) in 0.1M PB, and were followed by 2 h incubation at RT in a biotinylated secondary antibody goat anti-rabbit (1:1000; Jackson Laboratories) in 0.1M PB. After, sections were incubated in avidin biotin peroxidase complex (ABC Elite; Vector Laboratories; 60 min; RT), washed in 0.1M PB and developed with 3, 3-diaminobenzidine plus nickel (DAB; Vector Laboratories Kit; 2-5 min reaction). Sections were washed in 0.1M cacodylate buffer and postfixed with 1% osmium and 1.5% potassium ferrocyanide in cacodylate buffer for 1 hr at RT after sections were dehydrated in an ascending gradient of ethanol (ETOH; 35%, 50% 70%, 85% 90%). Sections were blocked-stained with 3% uranyl acetate in 70% ETOH for 2 hr at 4ºC before the 80% ETOH. Latest steps of dehydration were performed with 100% ETOH, propylene oxide followed by infiltration with epoxy resin (EMBed-812; Electron Microscopy Science, PA USA). Sections were flatted embedded between Aclar sheets and polymerized in an oven at 60ºC for 48 hrs. Selected areas of the cortex were trimmed and mounted on epoxy blocks and cut with a Leica EM UC7 ultramicrotome. Ultrathin sections (80 nm in thickness) were collected on single slot cupper grids with formvar. Ultrathin sections were observed with a JEOL-1400 transmission electron microscope (JEOL Ltd., Akishima Tokyo, Japan) and images were captured with an Orius™ SC200 CCD camera (Gatan Inc., Warrendale, PA, USA).

### Multiplex fluorescent in situ hybridization (FISH)

Mice were anesthetized and rapidly decapitated. The brain was quickly removed (< 2 min), placed into OCT, and flash frozen using 2-methylbutane chilled on dry ice. Tissue was kept on dry ice until cryosectioning. Cryosections (15 μm) were mounted directly onto Super Frost Plus slides, and fluorescence in situ hybridization (FISH) studies were performed according to the protocol for fresh-frozen sample using the RNAscope Multiplex Fluorescent v1 Assay (Advanced Cell Diagnostics, ACD, 320850). Probes (Advanced Cell Diagnostics) for Mm-Tacr1 (Cat. No. 428781), Mm-tdTomato (Cat. No. 317041) Mm-Nos1 (Cat. No. 437651), Mm-Chodl (Cat. No. 450211) and Mm-Sst (Cat. No. 404631) were hybridized for 2 h at 40°C in a humidified oven, followed by rinsing in wash buffer and a series of incubations to develop the hybridized probe signal. Human prefrontal cortex was cryosectioned at 20 μm and mounted directly onto Super Frost Plus slides and FISH studies were performed according to the protocol for fresh-frozen sample using the RNAscope Multiplex Fluorescent v2 Assay (ACD, 320850). Human probes from included Hu-TACR1 (Cat. No.17166A), Hu-NOS1 (Cat. No. 171594), Hu-CHODL (Cat. No.171634) and Hu-SST (Cat. No. 17145C). Sections were stained with DAPI (320858) and mounted with Prolong Diamond AntiFade (ThermoFisher, P36961).

### Stereotaxic injection and viruses

Animals were anaesthetized with 3–5% isoflurane and maintained at 1–2% isoflurane for the duration of the procedure. Animals were placed in a stereotaxic head frame (Kopf Instruments, Model 942 Small Animal Stereotaxic Instruments) and administered a subcutaneous dose of the analgesic (buprenorphine, 0.1 mg kg^−1^; ketoprofen, 5.0 mg kg^−1^) at the start of the procedure and was also administered ketoprofen BID for 48 h after the procedure. The scalp was shaved, local antiseptic applied (betadine), and a midline incision made to expose the cranium. The skull was aligned using cranial fissures. Local anaesthetic bupivacaine (2.5 mg kg^−1^) was topically applied on the skull at the site of drilling. A stainless-steel burr (Fine Science Tools, 19008-07) attached to a micro drill (Foredom Electric Co., K1070 High Speed Rotary Micromotor Kit, 2.35mm Collet) was used to create a burr hole. A custom pulled 3.5” glass capillary tube (Drummond, #3-00-203-G/X replacement) was loaded with AAV. Virus was infused at a rate of 2 nL/s using a microinjector (Drummond Scientific Company, Nanoinjector III, Cat.#3-000-207) *Tacr1*^*CreER*^ mice were unilaterally injected with 400 nl virus. The injection needle was left in place for an additional 5 minutes and then slowly withdrawn. Injections performed at the following coordinates for each brain region: S1BF: AP, −0.60 mm; ML, ± 2.9 mm; DV: 0.5 and 0.8 mm. The skin incision was closed using 2-3 simple interrupted sutures. A small amount (< 5 μl) of 3M Vetbond was placed on top of sutures to discourage grooming-related activities at the incision site. Mice were housed with original cage-mates and given 4 weeks to recover prior to experimentation.

The following viruses were used in this study: AAVr.EF1a.DIO.hChR2(H134R).EYFP-WPRE-HGH (Addgene 20298), AAV5-hSyn-Con/Foff-hChR2(h134R)-EYFP (UNC Vector Core) and AAV9-CaMKIIa-hChR2(H134R)-EYFP (Addgene 26969).

### Whole tissue clearing

Whole-tissue clearing on brains from AAVrg-DIO-ChR2-YFP (Addgene 20298) injected *Tacr1CreER* were using the CUBIC (unobstructed brain/body imaging cocktails) protocol previously described with minor modifications^22^. Briefly, animals were transcardially perfused with PBS then 4% PFA and brains and post-fixed overnight in 4% PFA at 4°C. Brains were incubated in 50% CUBIC R1 solution at 37°C on a nutating shaker until brains were translucent. Brains were then washed with IHC buffer incubated in primary antibody (GFP, 1:500 ThermoFisher A-11122), washes with IHC buffer and followed with the corresponding Alexa Fluor-conjugated secondary antibodies (1:500, ThermoFisher). Antibody incubations for completed for 1 week. Brains were refractive index matched using CUBIC R2 solution at 37°C on a nutating shaker until clear. Ribbon scanning microscopy of whole cortex was completed on an RS-G4 confocal microscope (Caliber ID, Andover, MD) fitted with a Nikon CFI90 20x, 1.00 NA, glycerol objective. Data were acquired at a resolution of ∼0.3 μm lateral and ∼10 μm axial. The image data was stitched and assembled using a 24-node compute cluster running custom software. Neurons were traced using the filaments tool in Imaris v9.7.2 (Bitplane) software first using the semi-automated Autopath tool then corrected using the manual editing tools.

### Circuit mapping

Subcellular ChR2-assisted circuit mapping (sCRACM) was performed as previously described^23^. We made whole-cell recordings from M1 neurons in coronal brain slices of mice aged P42-92 with thalamic or S1 axons expressing ChR2 and applied 1 ms flashes of blue light (∼ 1 mW) to evoke neurotransmitter release. To prevent polysynaptic activity, we further added TTX (1 mM) and 4-AP (100 mM). Data were acquired at 10 kHz using an Axopatch 700B (Molecular Devices) and Ephus software (www.ephus.org) on a custom-built laser scanning photostimulation microscope. Individual maps were repeated 2-4 times and averaged. Electrophysiology data were low pass filtered offline (1 kHz). Data analysis was performed with custom routines written in MATLAB (MathWorks).

### Chronic cranial windows

Two- to four-month-old mice underwent a craniotomy involving implantation of a sterile glass window and attachment of an aluminum chamber frame (Narishige Inc., CF-10). Mice were anaesthetized with 3– 5% isoflurane and maintained at 1–2% isoflurane for the duration of the craniotomy. The respiration rate and body temperature were continuously monitored throughout the procedure to ensure the appropriate level of anesthesia. A subcutaneous dose of analgesia, buprenorphine (0.1 mg kg^−1^) and ketoprofen (5.0 mg kg^−1^), was administered at the start of the procedure and twice a day for three additional days after the craniotomy. A local anesthetic bupivacaine (2.5 mg kg^−1^) was topically applied on skull at the site of the craniotomy. The craniotomy was centered over barrel cortex, approximately 1.5-2 mm posterior to bregma and 3 mm lateral to the midpoint between bregma and lambda. A custom cover glass consisting of a 4-mm round cover glass glued on a 5-mm round cover glass (CS-4R and CS-5R, Warner Instruments Inc.), and chamber frame were cemented (Lang Dental Manufacturing Company, Ortho-Jet™) onto the skull. Tamoxifen administration (75 mg kg^-1^) in *Tacr1*^*CreER*^ mice began 3 days post-surgery. During recovery the mice were acclimated to a custom treadmill for awake head-fixed data collection.

### Optogenetic Stimulation

Light stimulation and whisker stimulation experiments were performed in animals under awake head-fixed conditions. The light stimulus was delivered using a power-adjustable, TTL-controlled laser diode unit (CrystaLaser Inc., Reno, NV) connected to the optic fiber. The laser power at the tip of the fiber was set to 1 mW using a power meter (Melles Griot 13PM001, IDEX Inc., Rochester, NY). Air puffs were delivered using a pressure injector (Toohey Spritzer, Toohey Company, Fairfield, NJ) set to 30 psi. The light stimulation parameters were selected based on previous experiments (see Extended Data).^24, 25^ For activation of Channelrhodopsin, a 473-nm laser delivered 30 ms light pulse at 5 Hz for 1 s every 30 s. For activation archaerhodopsin (ArchT), 589-nm laser delivered 5 ms light pulses at 5 Hz for 1 s every 30 s. At least 10 stimulation trials were collected for each stimulation parameter set. For optogenetic silencing experiments, whisker stimulation was performed before (whisker_initial_) and after (whisker_final_) light inhibition. Additional experiments were conducted in a subset of mice to serve as control experiments. Control experiments were performed in either *Tacr1*^*CreER*^ mice (ChR2 negative) or *Tacr1*^*CreER*^ mice expressing eYFP or GCaMP6f rather than ChR2. Stimulation triggers and LDF data were recorded at 1 kHz (MP150, Biopac Systems Inc., Goleta, CA).

### Laser Doppler flowmetry

A laser Doppler flowmeter (LDF; Periflux 5000/411, Perimed AB, Jarfalla, Sweden) was used to acquire CBF data in response to optogenetic (light) or whisker stimulation. The LDF probe used has a tip diameter of 200 mm, operating wavelength of 780 nm and a sampling rate of 1 kHz. Time series spanning 30 s were obtained from all trials starting 5 s prior to stimulation onset. The LDF time series were low-pass filtered with a rectangular cut-off of 4Hz and down-sampled to 10 Hz. An air puffer was also placed in front of the contralateral whisker pad (50 ms puffs, delivered at 5 Hz). LDF was used to measure changes in cerebral blood flow (CBF). The change in CBF (ΔCBF) was calculated as (CBF_time_-CBF_baseline_) / CBF _baseline_ x 100. CBF _baseline_ was determined as the mean CBF during the 4 s before whisker or light stimulation. Time series from all trials (10 to 12 trials) were averaged and converted to percent change for each animal and then averaged across animals. The change in maximum (ΔCBF) was determined as the average value during 1 to 3 s after whisker or light stimulation. maximum CBF (%) difference was determined as the max. ΔCBF light ON – max. ΔCBF light OFF (ChR2) or ΔCBF light OFF – max. ΔCBF light ON (ArchT).

### Two-photon microscopy

In vivo two-photon microscopy cortical imaging was performed using a Ultima IV microscope (Bruker Nano, Inc.) coupled to an ultra-fast laser (Excite X3, Newport Spectra-Physics, Inc). Awake, head-fixed mice were placed on a custom-made setup designed to accommodate light and sensory stimulation during live brain imaging. Two-photon images were acquired using a 16x water immersion objective lens (0.80 NA, Nikon, Inc.) at a wavelength of 920 nm for studies imaging YFP/GFP, GCaMPf and vasculature. Images were obtained with a maximum field-of-view of 800 × 800 μm. We calibrate the laser power output through the objective lens and use <40 mW for time series imaging. Stimulus onset (t=0) and imaging recordings were synchronized using a National Instruments board that tracked the start-of-frame trigger.

### Vessel analysis

A subcutaneous injection of sulforhodamine 101 (SR 101, Thermo Fisher Scientific, 0.2 μl g^-1^) was administered in mice to visualize the cerebral vasculature in vivo. We imaged parenchymal arterioles and capillary branches from the middle cerebral artery. We identified a pial artery in the somatosensory cortex and traced its branching parenchymal arterioles and capillaries (cortex depth, approximately 0-500 μm). To distinguish arterioles and capillaries in vivo, selection of arterioles and capillaries were guided by eYFP labeling of VSMCs (*NG*^*CreER*^-GCaMP6f). Ten stimulation trials were acquired and averaged for each field of view. Three to seven locations were acquired per imaging session. Multiple imaging sessions were collected on separate days per mouse and arteriolar and capillary dilation responses were averaged across all sessions for each mouse. To determine percent change in diameter relative to baseline, the time series images were first filtered with a Gaussian filter and background subtracted with a rolling ball of 50 pixels. The diameter was measured as the full-width half-max of the vessel profile using custom Matlab routines. Time series were then smoothed using the default smooth function in MATLAB (Savitzky-Golay finite impulse response, 5 points). The change in diameter (ΔD/D) vessels was determined as (diameter_time_ - diameter_baseline_) /diameter_baseline_. Diameter_baseline_ was determined as the mean diameter during the 4 s before whisker or light stimulation. The maximum vessel dilation was determined as the maximum value before (t < −1 s) or after (t > 1.5 s) whisker or light stimulation (t = time). To determine latency onset to dilate, a line was fitted through 20% and 80% of the maximum value. The latency onset to dilate was considered the time difference between the x-intercept of the line and the start of the whisker or light stimulation.

### Calcium (Ca^2+^) imaging

For determination of calcium responses in GCaMP6f labeled neurons in *Tacr1*^*CreER*^-GCaMP6f mice, or VSMCs in *NG2*^*CreER*^:GCaMP6f mice, a region of interest (ROI) was selected encompassing individual cells or VSMCS. Ca^2+^ transients in regions of interested were longitudinally recorded by two-photon microscopy (excitation: 920 nm). Cell morphology, distinguishable by YFP labeling, was used to unambiguously identify pericytes (vs. VSMC or oligodendrocytes) in vivo. In addition, VSMC identity was confirmed post-hoc from Z-stacks of imaging sessions based on a combination of branching order, vessel diameter and distance from the parenchymal arteriole^19, 26^. Ca^2+^ responses were quantified with MATLAB. Briefly, the change in the Ca^2+^ signal (ΔF/F) was calculated as (fluorescence _time_-fluorescence _baseline_)/ fluorescence _baseline_. Fluorescence baseline intensity was determined as the average intensity for all time points for spontaneous activity measurements. For VSMCs, the minimum calcium response was determined as the minimum value after (t > 1.5 s) whisker or light stimulation. To determine latency onset to relaxation, a line was fitted through 20% and 80% of the maximum value. The latency onset to relaxation was considered the time difference between the x-intercept of the line and the start of the whisker or light stimulation. For neurons, the maximum calcium response was determined as the maximum value after (t > 1.5 s) whisker or light stimulation. Time series were smoothed using the customized smooth function in MATLAB (Savitzky-Golay finite impulse response, 5 points, for multiple vectors).

### Statistical Analyses

All statistical analyses were performed using Prism 9 (GraphPad Software) or MATLAB (R2019a or 2020). Two group comparisons were analyzed using a two-tailed Student’s *t*-test (paired or unpaired as indicated in figure legends) or non-parametric analyses. Multiple group comparisons were analyzed using a one-way ANOVA, followed by a post hoc Bonferroni analysis to correct for multiple comparisons. No data were excluded when performing statistical analysis. The s.e.m. was calculated for all experiments and displayed as errors bars in graphs. Statistical details for specific experiments (e.g., exact n values and what n represents, precision measures, statistical tests used and definitions of significance) can be found in figure legends. Values are expressed as mean ± s.e.m. No animals were excluded from analyses.

## Acknowledgements

We thank the donors and their loved ones for making our human studies possible and the Brain Tissue Donation Program at the University of Pittsburgh and the NIH NeuroBioBank for providing human prefrontal cortical sections. This work was funded in part by NIH grants F31-NS106724 to CFR, R01-NS090444 and R01-NS117515 to ALV, and R01-NS119410 to SER, as well as the Alzheimer’s Disease Research Center (ADRC) of the University of Pittsburgh.

## References

1. Attwell, D. et al. Glial and neuronal control of brain blood flow. Nature 468, 232–243 (2010).

2. Iadecola, C. The pathobiology of vascular dementia. Neuron 80, 844–866 (2013).

3. Iadecola, C. Neurovascular regulation in the normal brain and in Alzheimer’s disease. Nat Rev Neurosci 5, 347–360 (2004).

4. Girouard, H. & Iadecola, C. Neurovascular coupling in the normal brain and in hypertension, stroke, and Alzheimer disease. J Appl Physiol (1985) 100, 328–335 (2006).

5. Raichile, M.E. Behind the scences of funtional brain imaging: A historical and physiological perspective. Proc Natl Acad Sci USA 95, 765–772 (1998).

6. Iadecola, C. et al. Nitric oxide synthase-containing neural processes on large cerebral arteries and cerebral microvessels. Brain Res 606, 148–155 (1993).

7. Iadecola, C. Regulation of the cerebral microcirculation during neural activity: is nitric oxide the missing link? Trends Neurosci 16, 206–214 (1993).

8. Duchemin, S., Boily, M., Sadekova, N. & Girouard, H. The complex contribution of NOS interneurons in the physiology of cerebrovascular regulation. Front Neural Circuits 6, 51 (2012).

9. Tasic, B. et al. Adult mouse cortical cell taxonomy revealed by single cell transcriptomics. Nat Neurosci 19, 335–346 (2016).

10. Paul, A. et al. Transcriptional Architecture of Synaptic Communication Delineates GABAergic Neuron Identity. Cell 171, 522–539 e520 (2017).

11. Dittrich, L. et al. Cortical nNOS neurons co-express the NK1 receptor and are depolarized by Substance P in multiple mammalian species. Front Neural Circuits 6, 31 (2012).

12. Tamamaki, N. & Tomioka, R. Long-Range GABAergic Connections Distributed throughout the Neocortex and their Possible Function. Front Neurosci 4, 202 (2010).

13. Tomioka, R. et al. Demonstration of long-range GABAergic connections distributed throughout the mouse neocortex. European Journal of Neuroscience 21, 1587–1600 (2005).

14. Huang, H. et al. Generation of a NK1R-CreER knockin mouse strain to study cells involved in Neurokinin 1 Receptor signaling. Genesis 54, 593–601 (2016).

15. Fernandez-Klett, F., Offenhauser, N., Dirnagl, U., Priller, J. & Lindauer, U. Pericytes in capillaries are contractile in vivo, but arterioles mediate functional hyperemia in the mouse brain. Proc Natl Acad Sci U S A 107, 22290–22295 (2010).

16. Hall, C.N. et al. Capillary pericytes regulate cerebral blood flow in health and disease. Nature 508, 55–60 (2014).

17. Kisler, K. et al. Pericyte degeneration leads to neurovascular uncoupling and limits oxygen supply to brain. Nat Neurosci 20, 406–416 (2017).

18. Alarcon-Martinez, L. et al. Interpericyte tunnelling nanotubes regulate neurovascular coupling. Nature 585, 91–95 (2020).

19. Grant, R.I. et al. Organizational hierarchy and structural diversity of microvascular pericytes in adult mouse cortex. J Cereb Blood Flow Metab 39, 411–425 (2019).

20. Hartmann, D.A. et al. Pericyte structure and distribution in the cerebral cortex revealed by high-resolution imaging of transgenic mice. Neurophotonics 2, 041402 (2015).

21. Madisen, L. et al. A toolbox of Cre-dependent optogenetic transgenic mice for light-induced activation and silencing. Nat Neurosci 15, 793–802 (2012).

22. Muntifering, M. et al. Clearing for Deep Tissue Imaging. Curr Protoc Cytom 86, e38 (2018).

23. Hooks, B.M. et al. Organization of cortical and thalamic input to pyramidal neurons in mouse motor cortex. J Neurosci 33, 748–760 (2013).

24. Vazquez, A.L., Fukuda, M. & Kim, S.G. Inhibitory Neuron Activity Contributions to Hemodynamic Responses and Metabolic Load Examined Using an Inhibitory Optogenetic Mouse Model. Cereb Cortex 28, 4105–4119 (2018).

25. Vazquez, A.L., Fukuda, M., Crowley, J.C. & Kim, S.G. Neural and hemodynamic responses elicited by forelimb- and photo-stimulation in channelrhodopsin-2 mice: insights into the hemodynamic point spread function. Cereb Cortex 24, 2908–2919 (2014).

26. Rungta, R.L., Chaigneau, E., Osmanski, B.F. & Charpak, S. Vascular Compartmentalization of Functional Hyperemia from the Synapse to the Pia. Neuron 99, 362–375 e364 (2018).

