## Extended Data for "Long-range inhibitory neurons mediate cortical neurovascular coupling"

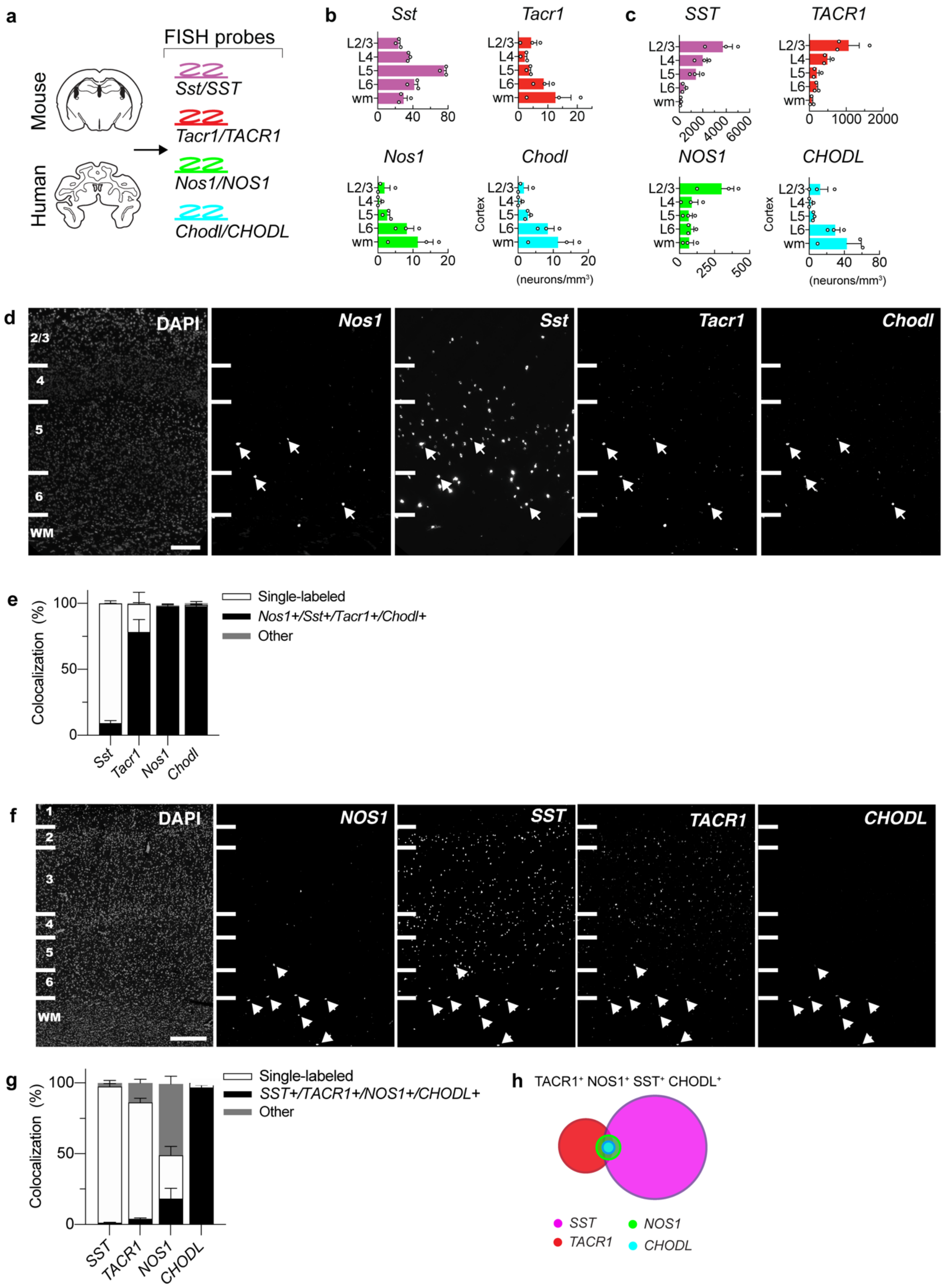

a
FISH probes
Mouse
Human
Sst/SST
Tacr1/TACR1
Nos1/NOS1
Chodl/CHODL
b
Sst
Tacr1
Nos1
Chodl
Cortex
(neurons/mm³)
c
SST
TACR1
NOS1
CHODL
Cortex
(neurons/mm³)
d
DAPI
Nos1
Sst
Tacr1
Chodl
WM
e
Colocalization (%)
Single-labeled
Nos1+/Sst+/Tacr1+/Chodl+
Other
f
DAPI
NOS1
SST
TACR1
CHODL
WM
g
Colocalization (%)
Single-labeled
SST+/TACR1+/NOS1+/CHODL+
Other
h
TACR1⁺ NOS1⁺ SST⁺ CHODL⁺
SST
TACR1
NOS1
CHODL

**Extended Data Figure 1 | Co-expression of four markers—Tacr1, Nos1, Sst, and Chodl— defines a putative cell type that appears conserved and shows a similar laminar distribution across species. a**, Schematic of multiplex fluorescent in situ hybridization (FISH) and mRNA probes used in mouse and human cortex. **b**, Laminar distribution of somatostatin (*Sst*, purple), tachykinin receptor-1 (*Tacr1*, red), neuronal nitric oxide synthase (*Nos1*, green) and chondrolectin (*Chodl*, blue) neuron populations in mouse somatosensory cortex. **c**, Laminar distribution of *SST* (purple), *TACR1* (red), *NOS1* (green) and *CHODL* (blue) neuron populations in human prefrontal cortex. **d**, Representative fluorescent images at low magnification of cortical section labeled for DAPI, *Nos1*, *Sst*, *Tacr1*, and *Chodl* mRNAs by multiplex FISH in mouse. Example of cortical neurons expressing *Nos1*, *Sst*, *Tacr1*, and *Chodl* in mouse (white arrows). Lines on the left demarcate the approximate laminar boundary, numbers indicate cortical layer. Scale bar (200 ґm) in DAPI image applies to all panels. **e**, Quantification of the percent of quadruple-labeled neurons (*Sst*$^{+}$/*Tacr1*$^{+}$/*Nos1*$^{+}$ /*Chodl*$^{+}$) that make up the *Sst*, *Tacr1*, *Nos1* or *Chodl* populations in mouse, which correspond to 9.10% ± 1.99, 78.26% ± 9.52, 97.71% ± 1.17, 97.79% ± 1.15, respectively (n = 3, C57BL/6 mice). **f**, Representative fluorescent images at low magnification of cortical section labeled for DAPI, *NOS1*, *SST*, *TACR1*, and *CHODL* mRNAs by multiplex FISH in human. Example of cortical neurons expressing *NOS1*, *SST*, *TACR1*, and *CHODL* in human (white arrows). Lines on the left demarcate the approximate laminar boundary, numbers indicate cortical layer. Scale bar (500 μm) in DAPI image applies to all panels. **g**, Quantification of the percent of quadruple-labeled neurons (*SST*$^{+}$/*TACR1*$^{+}$/*NOS1*$^{+}$*CHODL*$^{+}$) that comprise the SST, TACR1, NOS1 and CHODL populations in human which corresponds to 1.27% ± 0.33, 3.88% ± 0.63, 18.22% ± 7.34 and 96.70% ± 1.74, respectively (n = 3 human subjects). **h**, Venn diagram depicting the intersection of the neuron populations in human (colors correspond to those in a). WM, white matter. Data are mean ± s.e.m.

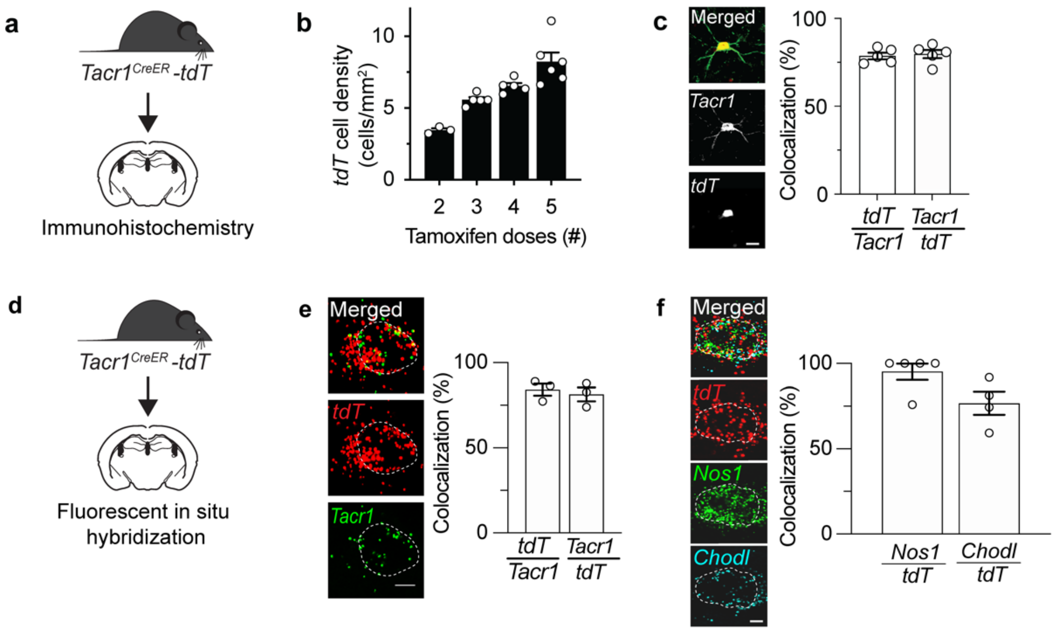


**Extended Data Figure 2 | *Tacr*$^{CreER}$-mediated tdTomato (*tdT*) expression is highly specific and efficient.** **a**, Immunohistochemistry was performed on tamoxifen administered *Tacr1*$^{CreER}$-tdT mice to evaluate the specificity and efficiency of *Tacr1*$^{CreER}$-mediated recombination in the cortex. **b**, *Tacr1*$^{CreER}$-mediated recombination is dependent on the number of tamoxifen doses administered (75 mg/kg SID). **c**, Representative IHC image and quantification of colocalization of Tacr1 immunoreactivity (green) and *Tacr1*$^{CreER}$-mediated *tdT* expression (red) in a cortical neuron (n = 5 mice) in tamoxifen-administered (five doses) *Tacr1*$^{CreER}$-*tdT* mice. Quantification of colocalization of endogenous tdT and Tacr1 protein within the cortex, demonstrated high specificity (78.7% ± 1.9) and recombination efficiency (79.8% ± 2.4). **d**, Fluorescent in situ hybridization (FISH) was performed on tamoxifen administered *Tacr1*$^{CreER}$-*tdT* mice to evaluate the specificity and efficiency of *Tacr1*$^{CreER}$-mediated recombination, and co-expression of *Nos1* and *Chodl* in mouse cortex. **e**, Representative multiplex FISH images of *Tacr1*$^{CreER}$-mediated *tdT* mRNA expression (red) colocalized with *Tacr1* mRNA (green) in a cortical neuron. Quantification of colocalization of *Tacr1* and *tdT* mRNA within the cortex, demonstrated high specificity (84.2% ± 3.5) and recombination efficiency (81.4% ± 4.1). **b**, Representative multiplex FISH images of *Tacr1*$^{CreER}$-mediated *tdT* mRNA expression (red) colocalized with *Nos1* and *Chodl* mRNA in a cortical neuron. Quantification of the colocalization of *tdT* mRNA with *Nos1* mRNA (green), or *Chodl* mRNA (blue) was 95.2% ± 4.8, and 76.7% ± 6.8 respectively (n = 4 mice). Scale bar, 5 μm. Data are mean ± s.e.m.

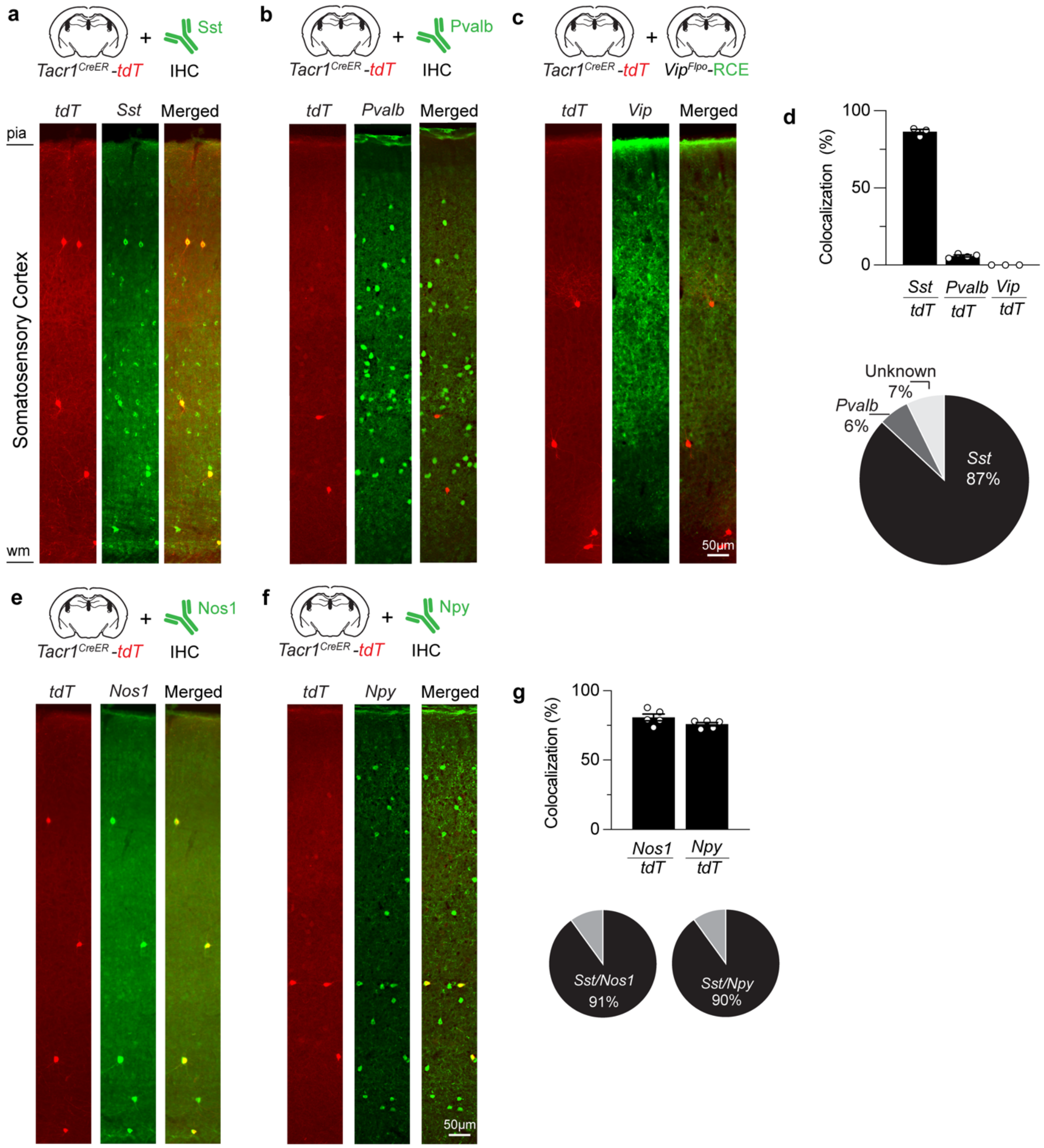


**Extended Data Figure 3 | tdTomato (*tdT*) expressing *Tacr1* neurons largely co-express *Sst*, *Nos1* and *Npy*. a-c**, Top: Schematic depicting immunohistochemistry approach for the indicated protein in tamoxifen-administered *Tacr1*$^{CreER}$-*tdT* mice to evaluate co-expression. To visualize VIP-expressing neurons, we used a genetic approach involving Vip-flp (JAX, no. 028578) and a derivative of the RCE:dual allele (Jax, no. 032036) that is only flp-responsive. Bottom: Representative low magnification images of immunostaining for somatostatin (*Sst*), parvalbumin (*Pvalb*), or vasointestinal peptide (*Vip*). **d**, Top: Quantification of Tacr1 *tdT*-expressing neurons with Sst, Pvalb, or Vip was 86.4% ± 1.6, 5.9% ± 0.7, and 0.0% respectively (n = 3 mice). Bottom: Venn diagram depicts that of three major inhibitory neurons evaluated (*Sst*, *Pvalb*, and *Vip*), *Tacr1* neurons belong to the *Sst* neuron population. **e-f**, Top: Immunohistochemistry for either neuronal nitric oxide synthase (*Nos1*) and neuropeptide Y (*Npy*) was performed in tamoxifen-administered *Tacr1*$^{CreER}$*-tdT* mice to evaluate co-expression. Bottom: Representative low magnification images of immunostaining for *Nos1* and *Npy*.; wm, white mater. **g**, Top: Quantification of colocalization of *tdT* with *Nos1* or *Npy* was 80.8% ± 2.48, and 75.8% ± 1.27 respectively (n = 3 mice). Bottom: Venn diagram depicts that within the *Sst* population, *Tacr1* neurons also co-expression *Nos1* and *Npy*. Data are mean ± s.e.m. Scale bar, 50 µm. DAPI, 4′,6-diamidino-2-phenylindole.

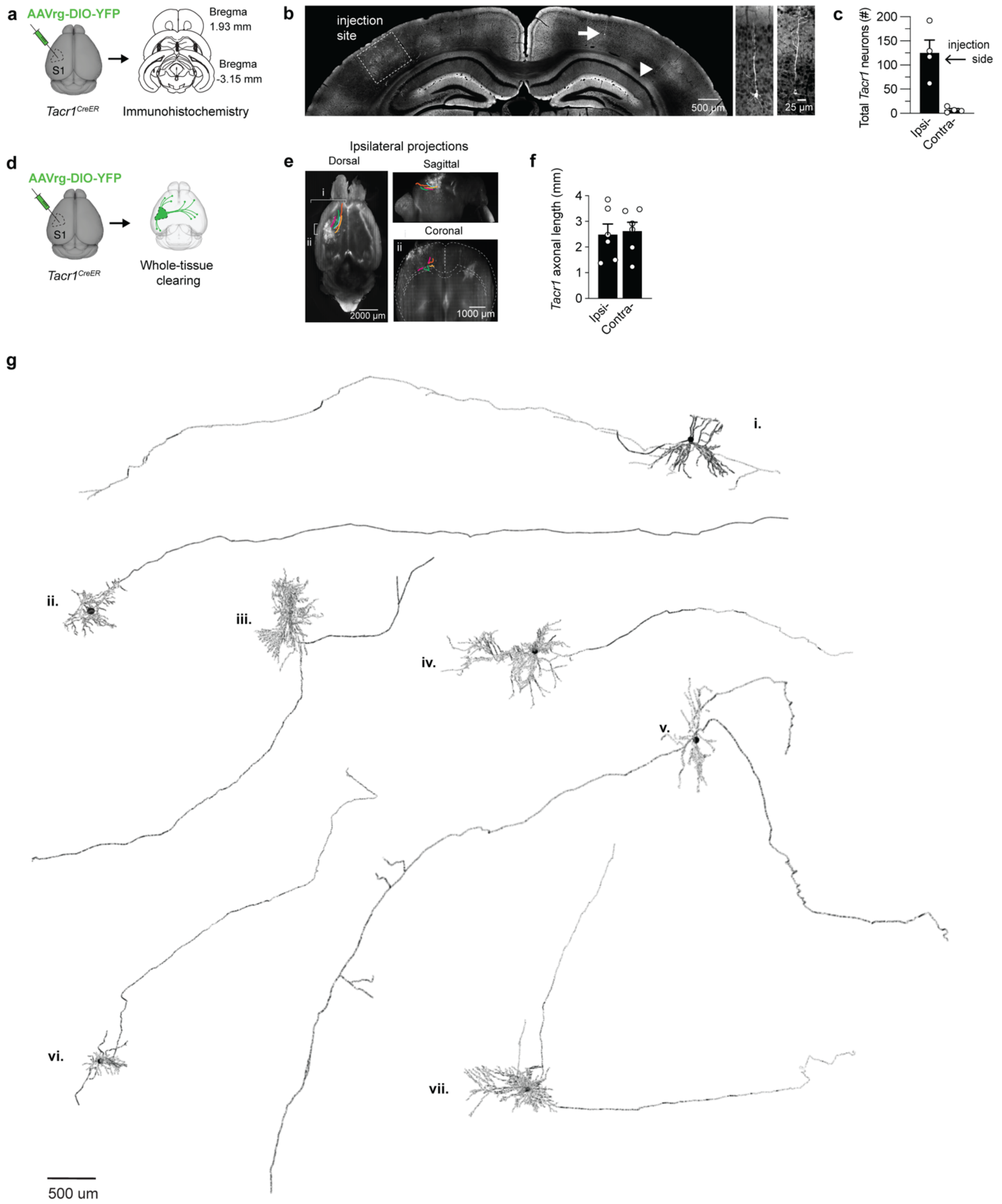


**Extended Data Figure 4 | Tacr1 morphology visualized using immunohistochemistry or whole-tissue clearing. a**, Schematic depicting experimental design. A Cre-dependent AAV-retrograde(rg) was injected into mouse somatosensory cortex and brains were processed for immunohistochemistry using antibodies against GFP/YFP to identify genetically-labeled Tacr1 neurons. **b**, Representative confocal microscopy image of YFP expression following the injection of AAVrg-YFP ino the S1 cortex of *Tacr1*$^{CreER}$ mice. Large scale coronal image shows the injection site (left) and contralateral retrogradely labeled *Tacr1* soma (right). Labeled Tacr1 soma are enlarged in images on the right (i, white arrow; ii, white arrow head). **c**, Quantification of retrogradely labeled Tacr1 soma on the ipsilateral and contralateral cortex. **d,** Schematic depicting experimental design. AAV-retrograde(rg) was injected into mouse somatosensory cortex and brains were processed for whole tissue clearing to trace and characterize Tacr1 neuron morphology. e, Ipsilateral projections of traced Tacr1 neurons in cleared brains (n = 2). Quantification of axonal length of Tacr1 neurons, which classifies Tacr1 neurons as long-range projection neurons. **g**, Representative traces of YFP-labeled Tacr1 neurons showing long-range axons and short, locally-branching dendrites.

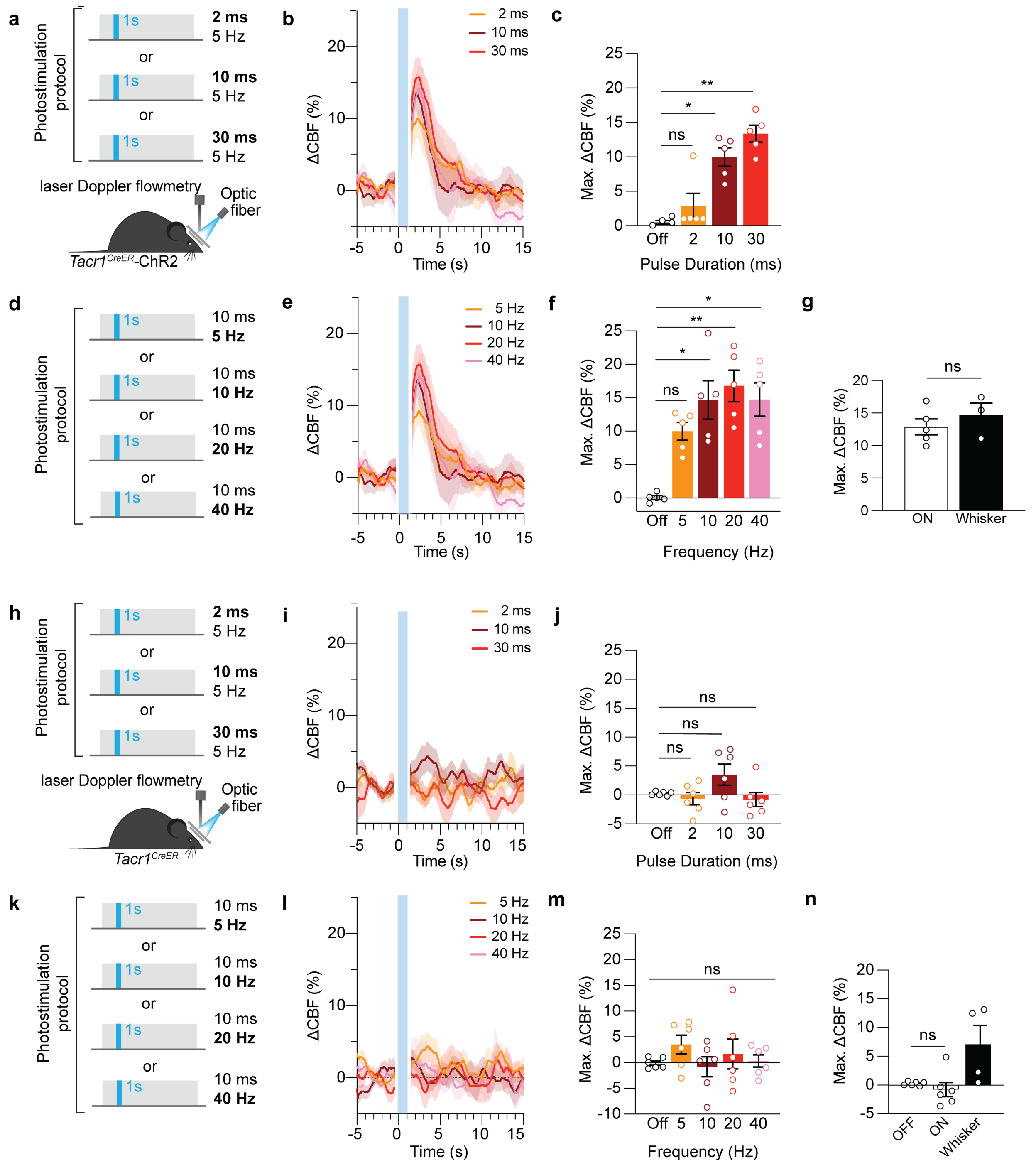


**Extended Data Figure 5 | Optical excitation is dose-dependent and does not elicit a hemodynamic response in ChR2 negative mice. a**, Optical excitation (1 mW, 1 s, 5 Hz) under varied pulse durations (2, 10, 30 ms) were evaluated in *Tacr1*$^{CreER}$-ChR2 mice. **b-c**, Optical excitation significantly increased CBF with increasing pulse duration (dose-dependent). Time course of the change in CBF (**a**) and maximum percent change in CBF (**b**) in *Tacr1*$^{CreER}$-ChR2 mice with varied pulse duration compared to pre-light baseline (mean CBF during 4 s prior to light stimulation). **d**, Optical excitation (1 mW, 1 s, 10 ms) under varied frequencies (5, 10, 20, 40 Hz) were evaluated in *Tacr1*$^{CreER}$-ChR2 mice. **e-f**, Optical excitation with increasing frequency significantly increased the CBF response in *Tacr1*$^{CreER}$-ChR2 mice at 10 and 20 Hz. Time course of the change in CBF (**e**) and maximum percent change in CBF (**f**) in ChR2-expressing mice with varied optogenetic frequency. **g**, Optical excitation (1 mW, 1 s, 5 Hz) under varied pulse durations (2, 10, 30 ms) were evaluated in ChR2 negative mice (*Tacr1*$^{CreER}$ alone mice). **h-i**, Optical excitation with varied pulse duration does not elicit a CBF response in ChR2 negative mice. Time course of the change in CBF (**e**) and maximum percent change in CBF (**f**) in ChR2 negative mice with varied pulse duration. **j**, Optical excitation (1 mW, 1 s, 10 ms) under varied frequencies (5, 10, 20, 40 Hz) were evaluated in ChR2 negative mice (*Tacr1*$^{CreER}$). **k-l**, Optical excitation with increasing frequency does not increase the CBF response in ChR2 negative mice (*Tacr1*$^{CreER}$). Time course of the change in CBF (**g**) and maximum percent change in CBF (**h**) in ChR2 negative mice with varied frequency. Statistical significance was determined by one-way ANOVA with a post hoc Dunnett's multiple comparison adjustment vs. control (pre-light baseline = mean CBF during 4 s prior to light stimulation). The vertical blue bars indicate the blue optogenetic stimulation period which often interfered with the laser Doppler readings of CBF, especially during light onset and offset. The data over this period is masked to exclude this artifact from the time course ΔCBF graphs. All data are mean ± s.e.m.

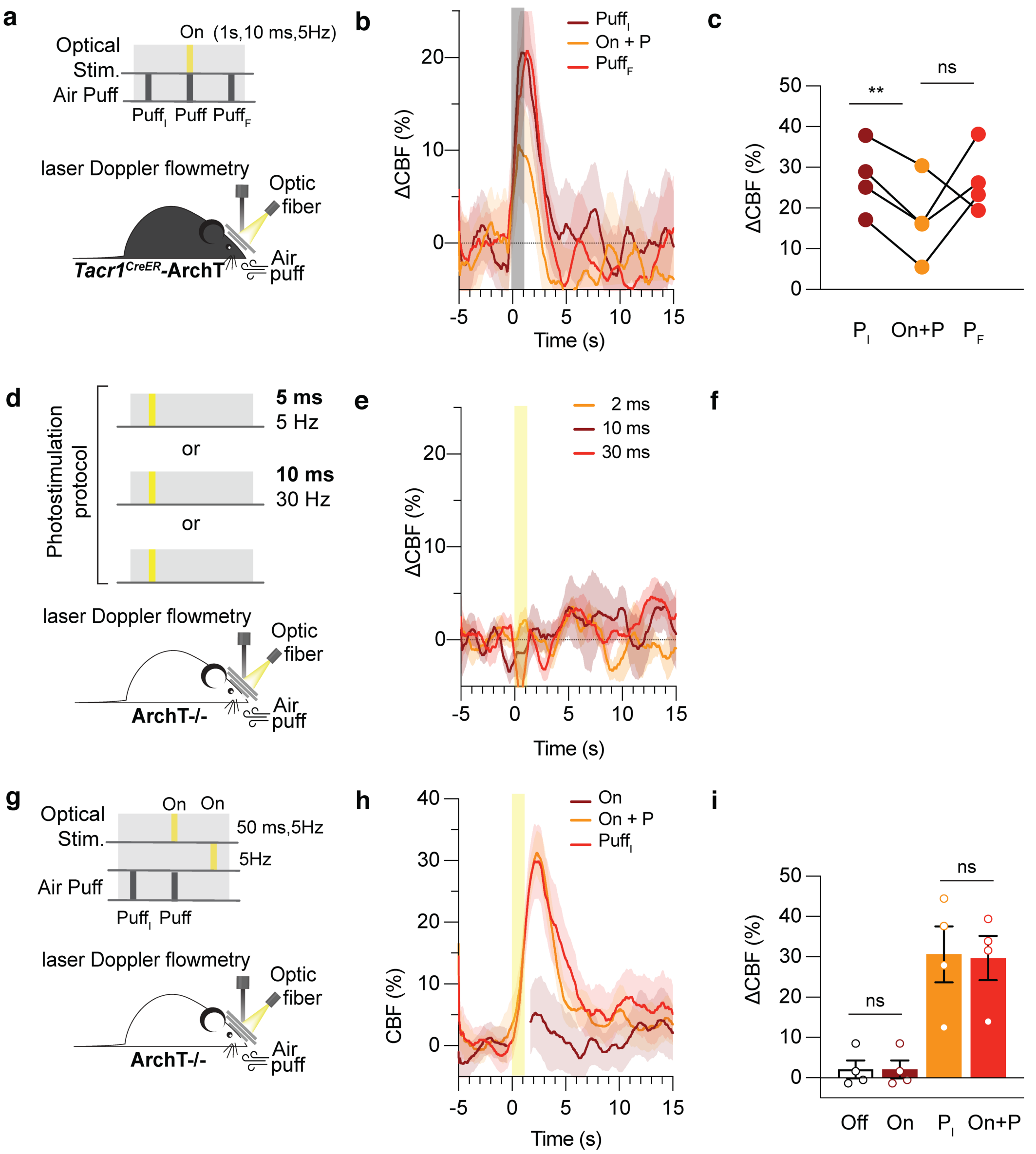


**Extended Data Figure 6 | Optogenetic inhibition (yellow light) does not elicit a change in CBF response, or inhibit whisker-evoked CBF response in mice lacking ArchT. a-c**, In *Tacr1*$^{CreER}$-ArchT mice, optogenetic inhibition with yellow light decreased the change in CBF relative to the initial sensory-evoked response ($P_I$, $^{**}P$ =0.0062) and the final sensory-evoked CBF response ($P_F$, $P$ = 0.416). **a**, Top: Optogenetic inhibition protocol (1 s, 5 Hz, 5 ms pulse width; yellow light) included 10 trials of an initial air puff ($P_I$), air puff and light (ON+P), and final air puff ($P_F$). Bottom: Experimental setup demonstrating continuous CBF recording by LDF during optical inhibition in the presence of sensory and/or light stimulation in awake, head-fixed mice. Time course of the change in CBF (**b**), maximum change in CBF (**c**) in mice expressing ArchT in Tacr1 neurons during sensory and/or light stimulation (n=4 mice, 10 trials per mouse). **d-f**, Optogenetic inhibition with yellow does not elicit a CBF response or inhibit the whisker-evoked CBF response mice lacking ArchT. **d**, Top: Optogenetic inhibition protocol (1 s, 5 Hz, 50 ms pulse width; yellow light) included 10 trials of interleaved initial air puff ($P_I$), air puff and light (ON+P), light (ON) and no light (OFF, 5 s prior to light stimulation). Bottom: Experimental setup demonstrating continuous CBF recording by LDF during optogenetic inhibition in the presence of sensory and/or light stimulation in awake, head-fixed mice. Time course of the change in CBF (**e**), maximum change in CBF (**f**) in mice lacking ArchT (OFF vs. ON, $P$ = 0.3910. Puff vs. ON+P, $P$ = 0.7994). All data are mean ± s.e.m (error bars and shaded areas). Statistical significance was determined by one-way ANOVA with a post hoc Dunnett's multiple comparison (**c**) or paired, parametric, two-tailed t-test (**f**). ns, not significant.

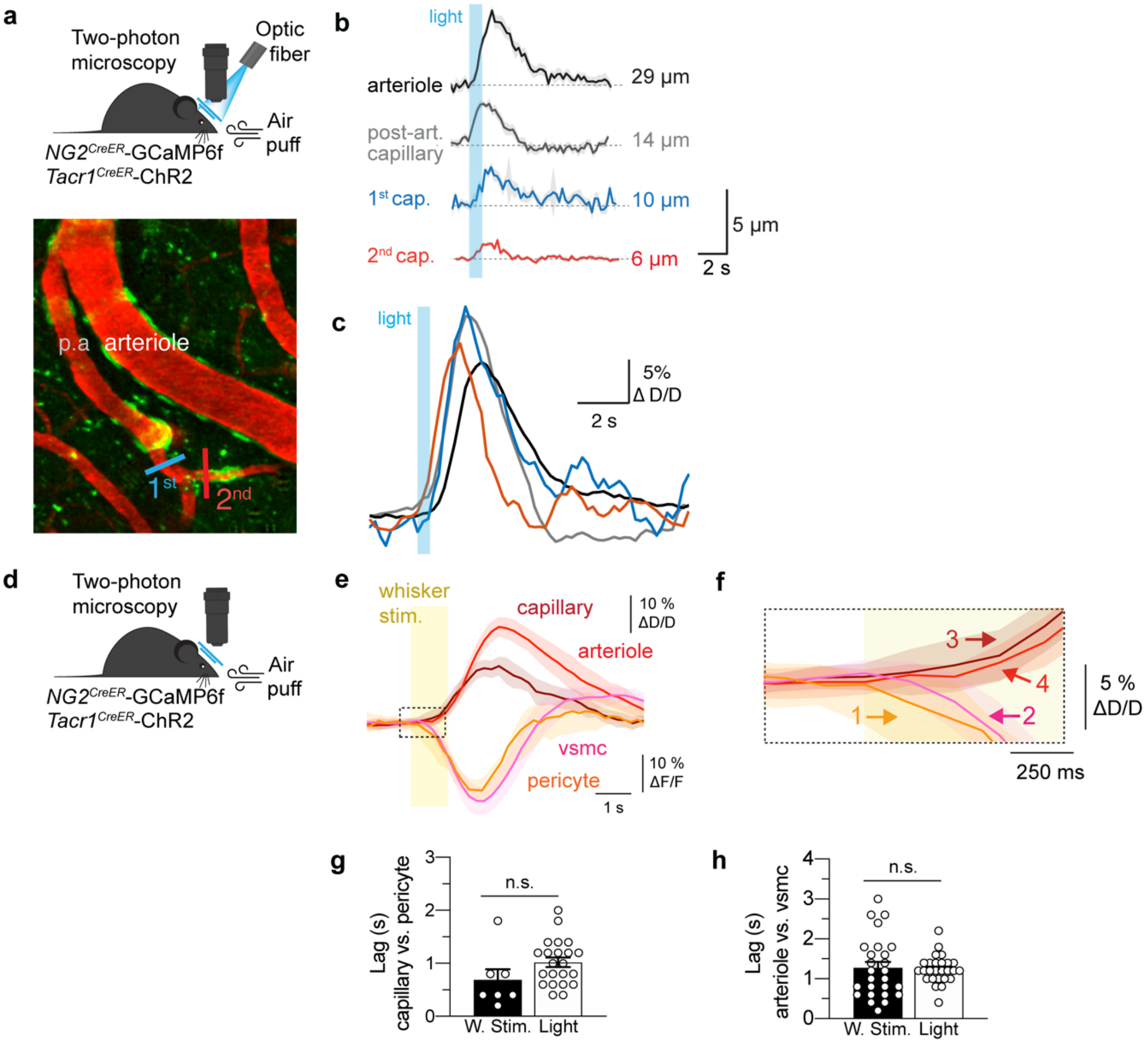


**Extended Data Figure 7 | Whisker-evoked CBF response has similar VSMCs/pericyte Ca$^{2+}$ signal and vascular dynamics to that observed upon activation of Tacr1 neurons. a**, Top: Schematic of optogenetic stimulation of Tacr1 neurons and simultaneous recording of Ca$^{2+}$ in VSMCs/pericyte and vessel diameter in triple transgenic mouse (*Tacr1*$^{CreER}$; *NG2* $^{CreER}$; *Rosa*$^{ls\text{-}lGCaMP6f}$) in which ChR2 is selectively expressed in Tacr1 neurons by virtue of a Cre-dependent AAV with a neuron-specific promoter (AAV9-Syn-DIO-ChR2). Bottom: *In vivo* two-photo microscopy image (maximum intensity projection) showing an example arteriole and its branching capillaries in mouse. **b**, Change in vessel diameter in vessels labeled in (**a**). Resting vessel diameter shown on right. **c**, Overlay of the CBF response of each vessel demonstrates that CBF onset occurs in 2$^{nd}$ order branching capillaries (red), followed by 1$^{st}$ order/post-arteriole capillaries (blue), finally the pial arteriole (black). A transient decrease in cytosolic free Ca$^{2+}$ signal in either pericytes and VSMCs, consistent with relaxation, accompanied by vasodilation of both capillaries and arterioles. **d-f**, Whisker-evoked CBF response has similar VSMCs/pericyte Ca$^{2+}$ signal and vascular dynamics. **d**, Schematic of sensory-evoked stimulation (air puff) and simultaneous recording of Ca$^{2+}$ in VSMCs/pericyte and vessel diameter in triple transgenic mouse (*Tacr1*$^{CreER}$:*NG2*:GCaMP6f). **e**, Time course of the change in VSMC/pericyte Ca$^{2+}$ signal and subsequent vessel diameter. Area outlined by black dashed rectangle shown in (f). **f**, VSMC/pericyte relaxation precedes vascular dilation during whisker stimulation. Numbers indicate response onset to whisker stimulation; colors correspond to the cell type shown above. **g-h**, Lag (seconds) between pericyte relaxation (F/F) and capillary dilation (D/D) (**j**) or VSMC relaxation (F/F) and arteriolar dilation (D/D) during light or whisker stimulation. **g**, There was no difference in the (lag) time between the onset of pericyte relaxation and capillary dilation during whisker stimulation compared *Tacr1*-evoked (light) stimulation. **h**, There was no difference in the (lag) time between the onset of VSMC relaxation and arteriolar dilation during whisker stimulation compared *Tacr1*-evoked (light) stimulation.

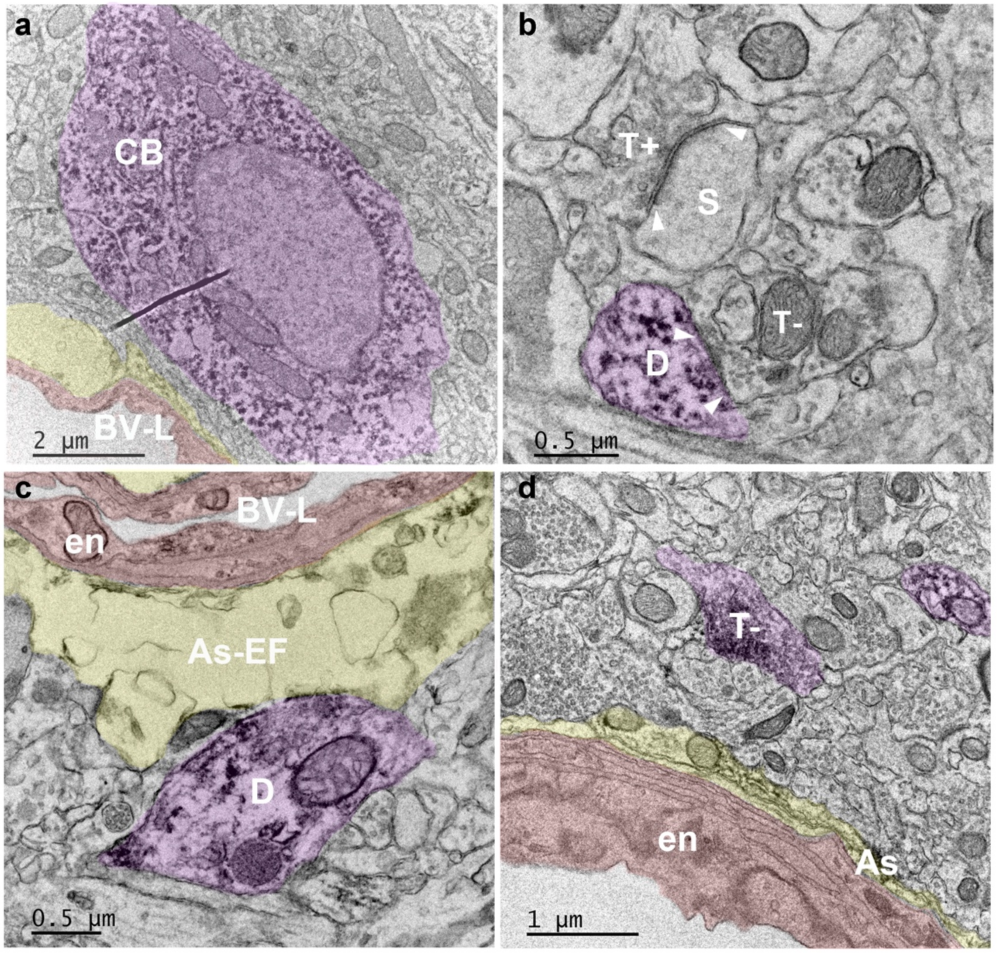


**Extended Data Figure 8 | Transmission electron microscopy of perivascular *Tacr1* neural processes.** Tacr1 neurons (cell bodies, dendritic and axonal processes) are false colored in purple. a, Example of a perivascular Tacr1 cell body (CB, purple). b, Tacr1 dendrite (D, purple) forming a symmetrical synapse (white arrows heads) with a putative inhibitory terminal (T-) that contains small and clear pleomorphic synaptic vesicles. Putative excitatory synaptic terminal (T+) forming an asymmetrical synapse (white arrow heads) with a spine (S). c, Tacr1 dendrite (D, purple) contacting an astrocytic endfoot (As-EF, yellow). d, Example of a perivascular Tacr1 presynaptic axonal terminal (T-, purple) in close proximity to a blood vessel. As: astrocyte; BV-L: blood vessel lumen; D: dendrite; en: endothelium; T-: putative inhibitory presynaptic terminal. T+: putative excitatory presynaptic terminal. (n = 2 mice).
